# Local brain-state dependency of effective connectivity: evidence from TMS–EEG

**DOI:** 10.1101/2021.10.01.462795

**Authors:** Ida Granö, Tuomas P. Mutanen, Aino E. Tervo, Jaakko O. Nieminen, Victor H. Souza, Matteo Fecchio, Mario Rosanova, Pantelis Lioumis, Risto J. Ilmoniemi

## Abstract

**Background:** Spontaneous cortical oscillations have been shown to modulate cortical responses to transcranial magnetic stimulation (TMS). If not controlled for, they might increase variability in responses and mask meaningful changes in the signals of interest when studying the brain with TMS combined with electroencephalography (TMS–EEG). To address this challenge in future closed-loop stimulation paradigms, we need to understand how spontaneous oscillations affect TMS-evoked responses.

**Objective:** To describe the effect of the pre-stimulus phase of cortical mu (8–13 Hz) and beta (13–30 Hz) oscillations on TMS-induced effective connectivity patterns.

**Methods:** We applied TMS to the left primary motor cortex and right pre-supplementary motor area of three subjects while recording EEG. We classified trials off-line into positive- and negative-phase classes according to the mu and beta rhythms. We calculated differences in the global mean-field amplitude (GMFA) and compared the cortical spreading of the TMS-evoked activity between the two classes.

**Results:** Phase had significant effects on the GMFA in 11 out of 12 datasets (3 subjects × 2 stimulation sites × 2 frequency bands). Seven of the datasets showed significant differences in the time range 15–50 ms, nine in 50–150 ms, and eight after 150 ms post-stimulus. Source estimates showed complex spatial differences between the classes in the cortical spreading of the TMS-evoked activity.

**Conclusions:** TMS-evoked effective connectivity appears to depend on the phase of local cortical oscillations at the stimulated site. This may be crucial for efficient design of future brain-state-dependent and closed-loop stimulation paradigms.

## Introduction

Brain state affects cortical responses to transcranial magnetic stimulation (TMS; [1, 2, 3, 4, 5, 6, 7, 8, 9, 10]), which can be observed by combining TMS with electroencephalography (TMS–EEG). For instance, during different sleep stages or deep sedation, TMS–EEG reveals different effective connectivity patterns that show how the signal propagates from the stimulation site to other brain areas [1, 4]. However, the relationship between electrophysiological indices and effective connectivity has not been studied in depth. Noting that EEG signals provide a measure of brain state (projection of post-synaptic currents; [11]), we focus on the phase of these oscillatory signals as a reflection of the local brain state and its impact on effective connectivity patterns.

Any brain disorder involves altered brain states, which in turn may reflect aberrations in effective connectivity [12, 13, 14, 15, 16, 17] that work both as biomarkers [18, 19, 20, 21, 22, 23] and as targets for treatment [19]. TMS–EEG can provide stable markers of effective connectivity in health and disease (for a review, see [24]). However, pre-stimulus oscillations can modulate the cortical responses to TMS [3, 10, 25, 26]. Thus, if not experimentally controlled for, within-subject variability may mask meaningful changes of reactivity and measures of connectivity [27], compromising the sensitivity and specificity of TMS–EEG-derived measures as biomarker [28, 29, 30]. To address these challenges, there are ongoing efforts to develop brain-state-dependent and closed-loop stimulation paradigms [31, 32, 33, 34, 35, 36, 37, 38, 39, 40, 41]. Nevertheless, to recognize optimal oscillatory states for closed-loop stimulation, we must understand the basic mechanisms through which oscillations modulate cortical effective connectivity.

In the frontal lobe, both mu and beta rhythms (8–13 Hz, 13–30 Hz, respectively) can modulate corticospinal responses [25, 42, 43]. The sensorimotor mu rhythm has been demonstrated to play a role in modulating excitability [44] and inducing cortical plastic changes [10, 45]. In this work, we investigate the role of the phase of these two rhythms (mu and beta) in influencing effective connectivity when stimulating the left primary motor cortex (M1) and the right pre-supplementary motor areas (pre-SMA). As an indicator of effective connectivity, we use causal signal transfer from one cortical area to another and investigate it in the light of TMS-induced signal propagation, i.e., the pattern of TMS-evoked activity that spreads across the cortex. We hypothesize that any changes in effective connectivity patterns could occur through either of the following mechanisms or their combination:

a. The phase of the ongoing oscillation determines the local excitability state during TMS by modulating the efficacy of stimulation and the overall amplitude of TMS-evoked potentials (TEPs) with no effects on cortical effective connectivity.
b. The local excitability state determines which pathways are activated by the TMS pulse. This could lead to complex pattern changes that modulate TEPs both at the stimulated site and at connected regions. The efficacy of neural input is known to be modulated by the excitability of the receiving population [46]—this mechanism could play a role in determining which neurons are most affected by the TMS pulse.

In this study, we show new evidence about the role of oscillatory activity for signal propagation and effective connectivity within the human cortical circuits.

## Methods

### Data acquisition

Three healthy right-handed volunteer subjects (S1, female, 28 years old; S2, male, 41; S3, male, 43), were recruited. The Coordinating Ethics Committee of Helsinki University Hospital approved the study, and all subjects signed a written informed consent. During the experiment, the subject sat in a comfortable chair, fixating a cross 3 m away. To attenuate the perception of the click sound produced by the TMS pulse, the subject wore earmuffs [4, 47, 48, 49]. The remaining sound was masked by white noise combined with random bursts of recorded TMS click sounds [50] continuosly playing through in-ear earphones worn underneath the earmuffs. No subject reported hearing the coil click.

Biphasic TMS pulses were delivered through a figure-of-eight coil (70-mm radius; Nexstim Cooled Coil, Nexstim Plc, Finland) connected to a Nexstim NBS 4.3 eXimia stimulator. Coil positioning was guided by neuronavigation software (Nexstim) based on the individual’s T1-weighted magnetic resonance image (MRI) with electric-field visualization. EEG signals were recorded with 60 Ag/AgCl-sintered electrodes and a TMS-compatible sample-and-hold amplifier ([51]; eXimia EEG, Nexstim), bandpass-filtered at 0.1–350 Hz, and sampled at the rate of 1450 Hz. The scalp under the electrodes was scraped with conductive abrasive paste (OneStep AbrasivPlus, H + H Medical Devices, Germany) before the electrodes were filled with conductive gel (Electro-Gel, ECI, Netherlands). The impedance of each electrode was kept below 5 kΩ. The reference electrode was placed on the right mastoid and the ground on the right zygomatic bone. Motor evoked potentials (MEPs) were recorded with a Nexstim electromyography (EMG) system. The EMG electrodes were fixed in a belly–tendon montage on the right abductor pollicis brevis (APB) muscle. The coil location and orientation producing the largest APB muscle response (cortical representation of APB; [52, 53]) and the resting motor threshold (RMT) were determined for each subject before the TMS–EEG experiment. RMT was defined as the stimulation intensity that evoked MEPs with a minimum amplitude of 50 µV in a resting muscle in 5 out of 10 stimuli given to the optimal position [54].

Single-pulse TMS was applied to the left M1 at the cortical representation of APB and the right pre-SMA (Fig. 1). When stimulating M1, we rotated the coil to minimize both peripheral and scalp muscle activations to avoid contamination of the EEG by the sensory re-afferent proprioceptive feedback [55]. At the same time, we ensured that the early (< 30 ms) TEPs had an artifact-free peak-to-peak amplitude of 6–10 μV in the average reference, as proposed by [56, 57], by keeping the TMS intensity at approximately 90% of RMT [55]. This resulted in stimulation intensities of 60 V/m (calculated electric field strength with a locally fitted spherical head model at the stimulation target) for S1, 55 V/m for S2, 90 V/m for S3.

**Fig. 1.**
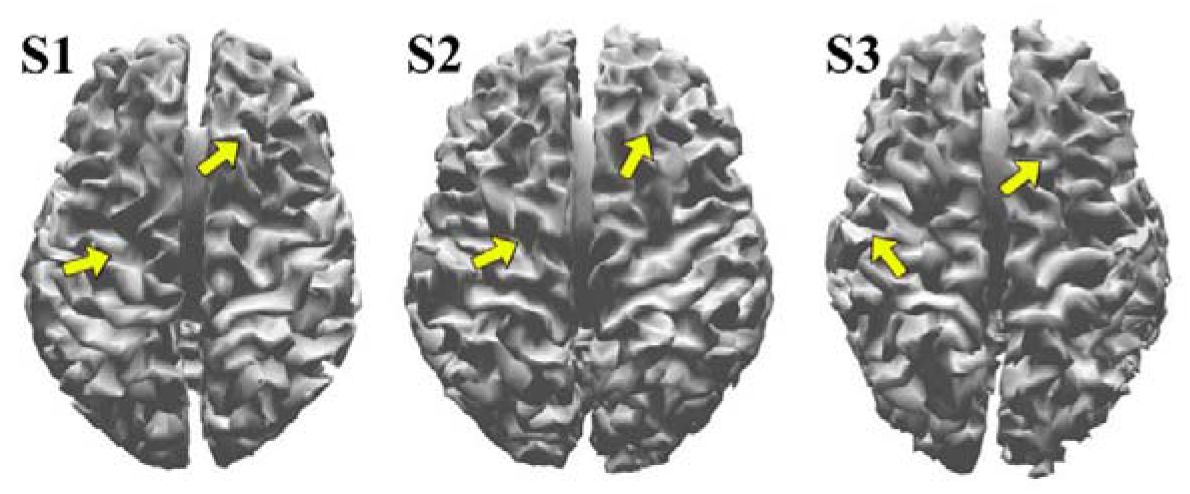
The approximate location and orientation of the maximum electric field: left M1 and right pre-SMA.

For the pre-SMA, we identified the rough stimulation area by individual anatomical landmarks as described by [49] and [58]. The final target and stimulation intensity were determined by the early TEP responses (6–10 μV), as proposed earlier [4, 49, 56, 57]. The final stimulation intensities at pre-SMA for S1, S2, and S3 were 100, 80, and 125 V/m, respectively. The stimuli were given at random interstimulus intervals of 2–2.3 s; 250 pulses were delivered to each target. A graphical user interface was utilized for evaluation of the TEPs during the experiment [57].

### Pre-processing

Data were pre-processed with custom-made MATLAB 2019a scripts [59] based on the EEGLAB toolbox [60]. The data were first filtered at 1–45 Hz with a 3^rd^-order zero-phase-shift Butterworth bandpass filter. Then, epochs were extracted with a time window of −1 to 1 s with respect to the TMS pulse. After visual inspection, we removed trials heavily contaminated by eye blinks or scalp-muscle activations. Then, data were re-referenced to the average potential and baseline corrected by subtracting the baseline average (−1000…−2 ms). Next, independent component analysis (ICA) separated the data into predominantly artefactual and neuronal components. These components were visually inspected for every trial; trials with highly distorted components were rejected. If trials were rejected based on independent components, ICA was computed again on the remaining data (number of remaining trials, after both trial-rejection steps: (mean ± std 233 ± 10.5, range 218–244). Independent components generated by eye blinks, eye movements, continuous muscle artifacts and electrode movement noise were removed from the data (mean ± std: 12 ± 2 components were removed per dataset).

### Phase evaluation

The trials were split manually into positive- and negative-phase classes, separately for mu and beta bands, based on the pre-stimulus phase of the frequency band in each trial. To help with decision-making, both raw and 4^th^-order Butterworth bandpass-filtered signals at the frequencies of interest were displayed from channel C3 (when stimulating M1) or F2 (when stimulating pre-SMA). A trial was classified as positive- or negative-phase if the filtered signal was near a positive or negative peak (less than 40° from peak) at the time of the TMS pulse, respectively, and the unfiltered signal was qualitatively similar in waveform to the filtered one (Fig. 2). This classification was done separately for each stimulation site and subject. This resulted in 72.6 ± 20.5 (mean ± std) trials in each class and a total of 12 datasets (2 stimulus locations × 3 subjects × 2 frequency bands), which were analysed separately for the effect of instantaneous phase at the stimulation onset.

**Fig. 2.**
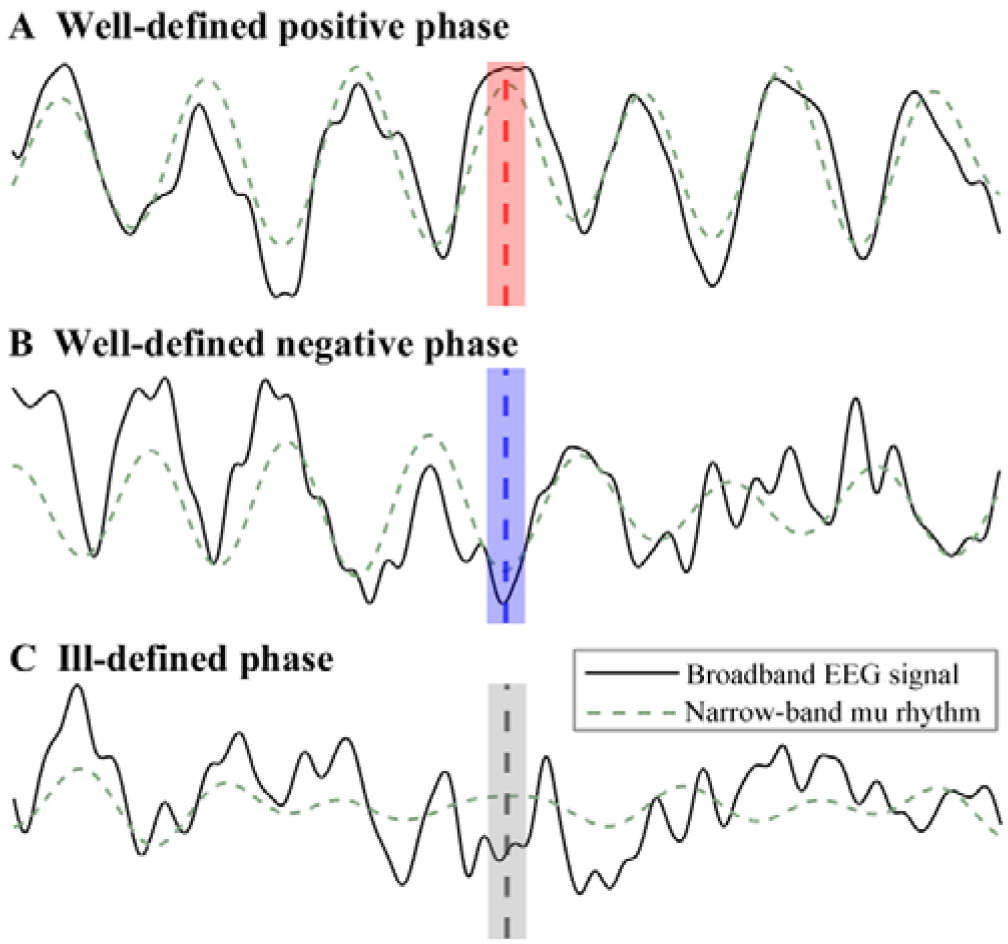
Sorting of trials into classes based on whether the TMS pulse was given near the peak or trough of the oscillation. A. A trial with a well-defined mu rhythm, evident as high similarity between the narrow-band and broadband EEG signals; the phase can be determined reliably and the trial classified as positive-phase. B. A trial with a well-defined mu rhythm, classified as negative-phase. The coloured bars illustrate the range within which the local maximum or minimum of a classified signal needs to be, corresponding to a maximum deviation of approximately 40° from 180° (negative phase) or 0° (positive phase). C. An ill-defined mu rhythm. At the vertical dashed line, the narrow-band signal suggests a positive phase classification; however, this is not evident from the broadband EEG signal. This trial would be left out of both classes.

### Correction of background oscillatory activity

When averaging evoked responses across trials, the usual assumption is that any background oscillations unrelated to the stimulus are attenuated by the averaging process. However, in trials classified according to the pre-stimulus phase, such background neural oscillations are consistent across trials and are consequently present in the averaged signal. This effect is of the opposite sign between the two classes, and if not properly addressed, may lead to incorrect interpretations. We assume that the averaged TEP in our case can be separated into two parts, the actual mean TEP and a waveform corresponding to the phase classification effect:

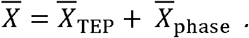

Thus,

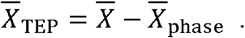

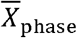 can be isolated from resting-state EEG, divided into trials, and sorted according to phase with the same technique as the real trials. When averaging these classified non-stimulated trials, all activity is cancelled out except for the phase effect. If one assumes that the TMS pulse does not interact with the background oscillation, this phase effect can be subtracted from each of the stimulated trials.

We applied this approach by extracting the pre-stimulus time period (−1000…0 ms) of each trial to use as non-stimulated trials with a time range of −500…500 ms. To match this length, we cut the stimulated trials to −500…500 ms when applying the correction.

The subtraction approach has been used previously, both with [26, 61] and without [62] TMS, to address effects on the mean response by classification according to phase. The authors of those studies reported that this correction of background oscillatory activity successfully removed the baseline differences originating from the phase classification, in line with our observations (Fig. 3).

**Fig. 3.**
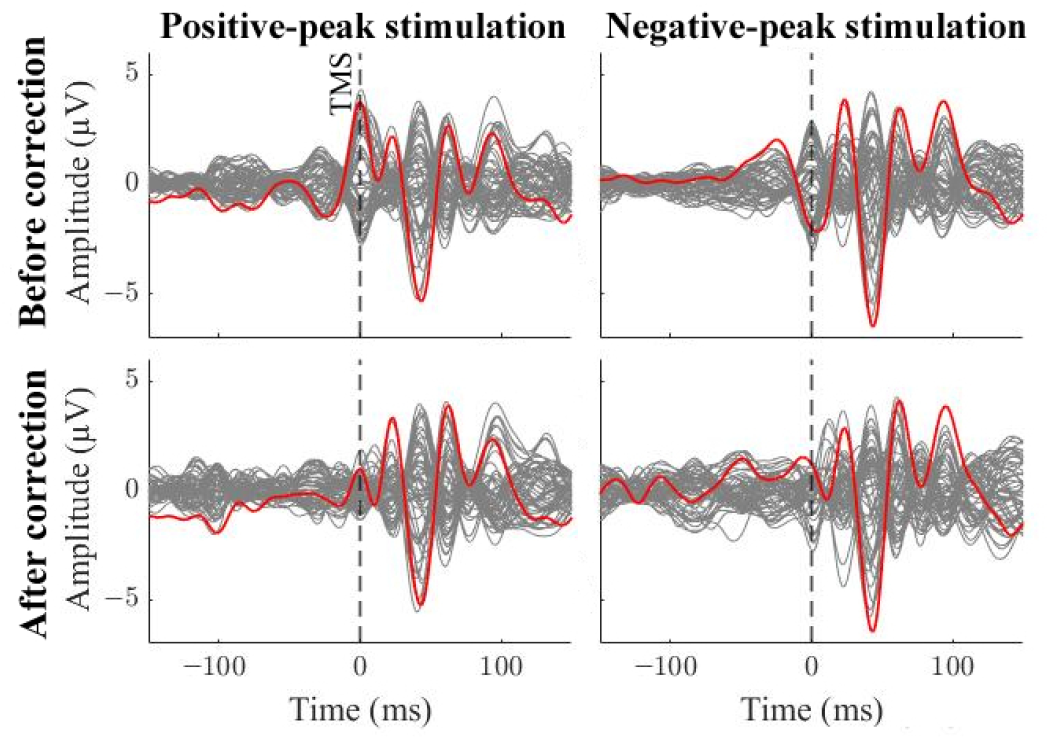
TEPs (S3, stimulation of pre-SMA) before and after the phase correction, after classification according to the beta rhythm. The traces show all channels (grey); the channel closest to the stimulation site (F2) is highlighted in red. The correction suppresses the large phase-related deflection around the TMS pulse.

### Statistics and source estimation

For each dataset, the global mean-field amplitude (GMFA, the square root of the sum of squared amplitudes at a given time, [63, 64]) was compared between the two classes. A threshold was set by taking the 95th percentile of the difference in GMFA of the pre-stimulus time period (−500…0 ms); any differences larger than this in the post-TMS time period (0…300 ms) were considered significant (see similar thresholding by [4]). For all significant time intervals, the mean EEG response over the time window was calculated for both classes separately to be utilized in the source estimation. Tikhonov-regularized minimum-norm estimates (MNE) of the sources were used for locating spatial differences of cortical activity between the two classes in these time intervals. The obtained MNE maps were thresholded for visualization to show only the cortical area corresponding to at least 60% of the maximum MNE amplitude.

For source estimation, the scalp, skull, and white-matter surfaces were extracted from the MRIs using the SimNIBS software [65]. The meshes were imported to MATLAB, decimated to ∼10,000 nodes, and cleaned from isolated surfaces, intersections, duplicate nodes, holes and inverted triangle normals using the iso2mesh package [66]. The lead fields were calculated with the boundary element method [67] assuming conductivity values 0.33, 0.0033 and 0.33 S/m for the intra-cranial cavity, skull and scalp, respectively. A Tikhonov-regularized MNE was used for the inverse calculations [68] with a regularization parameter of 0.1. Dipoles were placed normal to the white matter surface.

## Results

### TEPs

TEPs (Fig. 4) had waveforms with characteristic frequencies for each stimulation site similar to what has been reported previously [49, 55, 69]. Significant differences in GMFAs between the positive- and negative-phase classes were found in 11 out of 12 comparisons (Figs. 5 and 6). Seven of the datasets showed significant differences in the time range 15–50 ms, nine in 50–150 ms, and eight after 150 ms post-stimulus. The significant differences, as revealed by the source estimates, showed the most abundant differences in frontal and central regions, although differences were also found in parietal, temporal and occipital areas. We observed large inter-individual variability in both the times and locations of significant differences between the two classes.

**Fig. 4.**
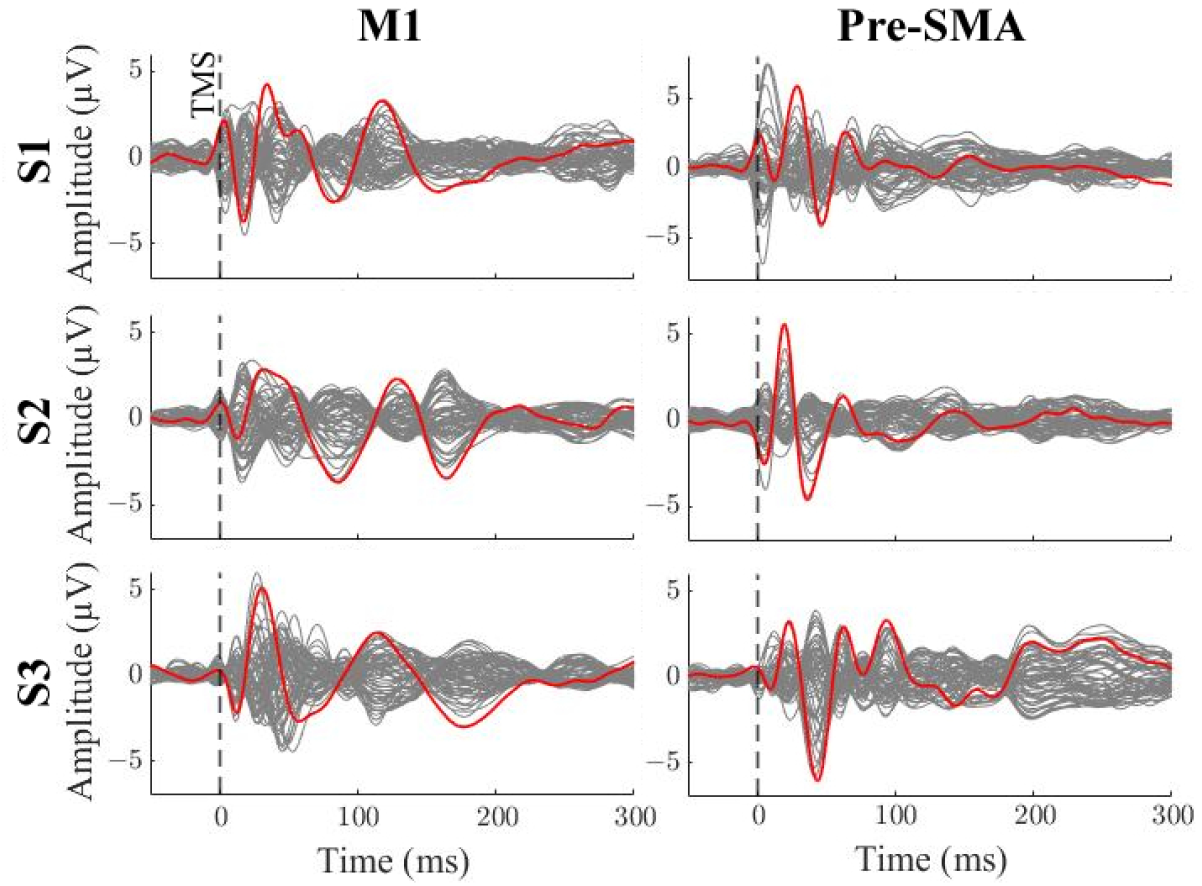
TEPs recorded at all EEG channels (grey traces) after stimulation of M1 and pre-SMA, all accepted trials included. The signal from the channel closest to the stimulation site (C3/F2) is highlighted in red.

**Fig. 5.**
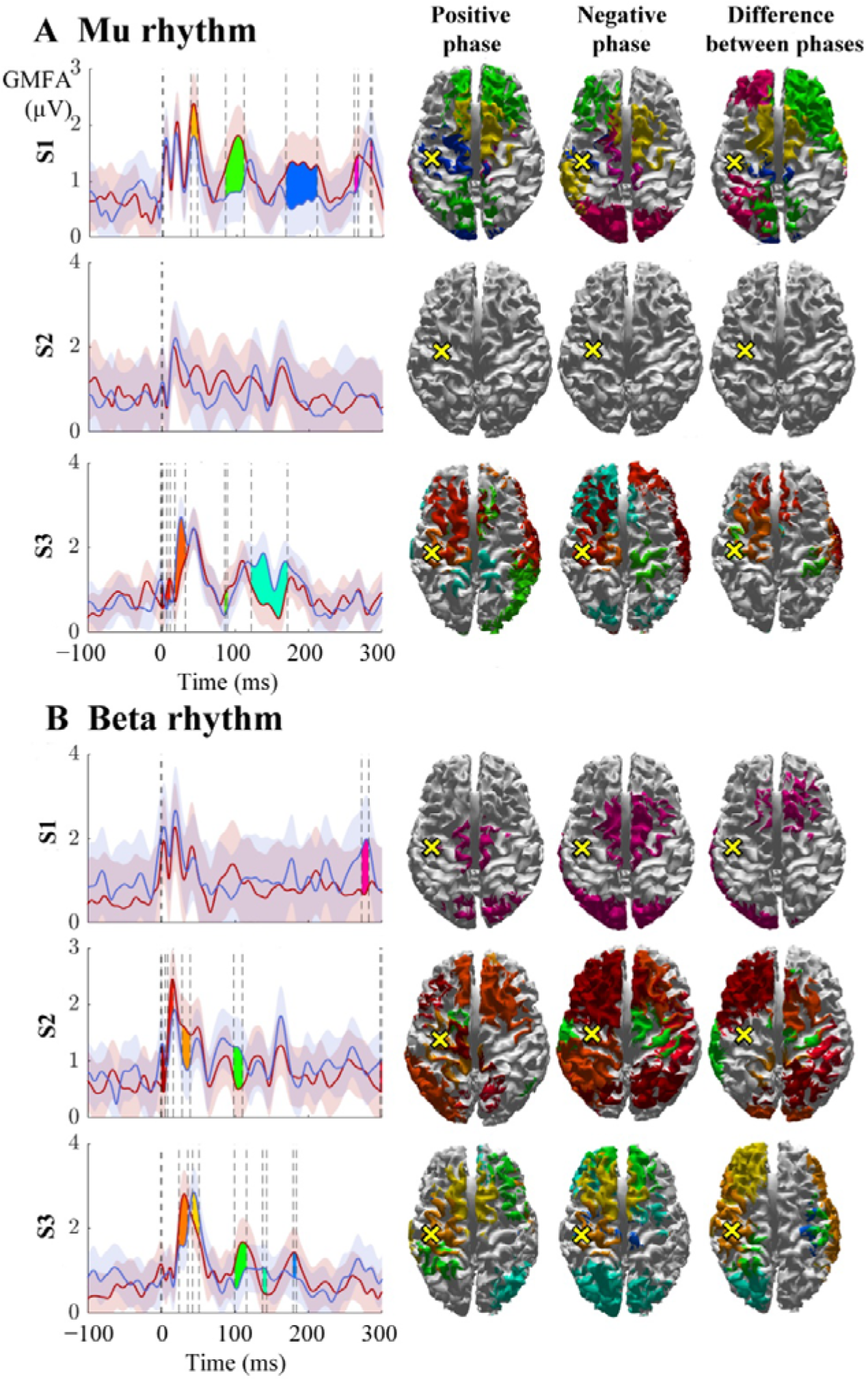
Global mean-field amplitudes (GMFA) of the positive-phase (red) and negative-phase (blue) conditions of mu (A) and beta (B) rhythms and their source estimates. The shaded areas indicate the GMFA ± the threshold for significance. Significant time intervals are marked with different colours. For each time interval, the corresponding time-averaged source estimates are shown on the right in the same colour. For each time interval, only sources stronger than 60% of the maximum amplitude are shown. The dark dashed vertical line indicates the time of the TMS pulse. The cross marks the stimulation location on M1.

**Fig. 6.**
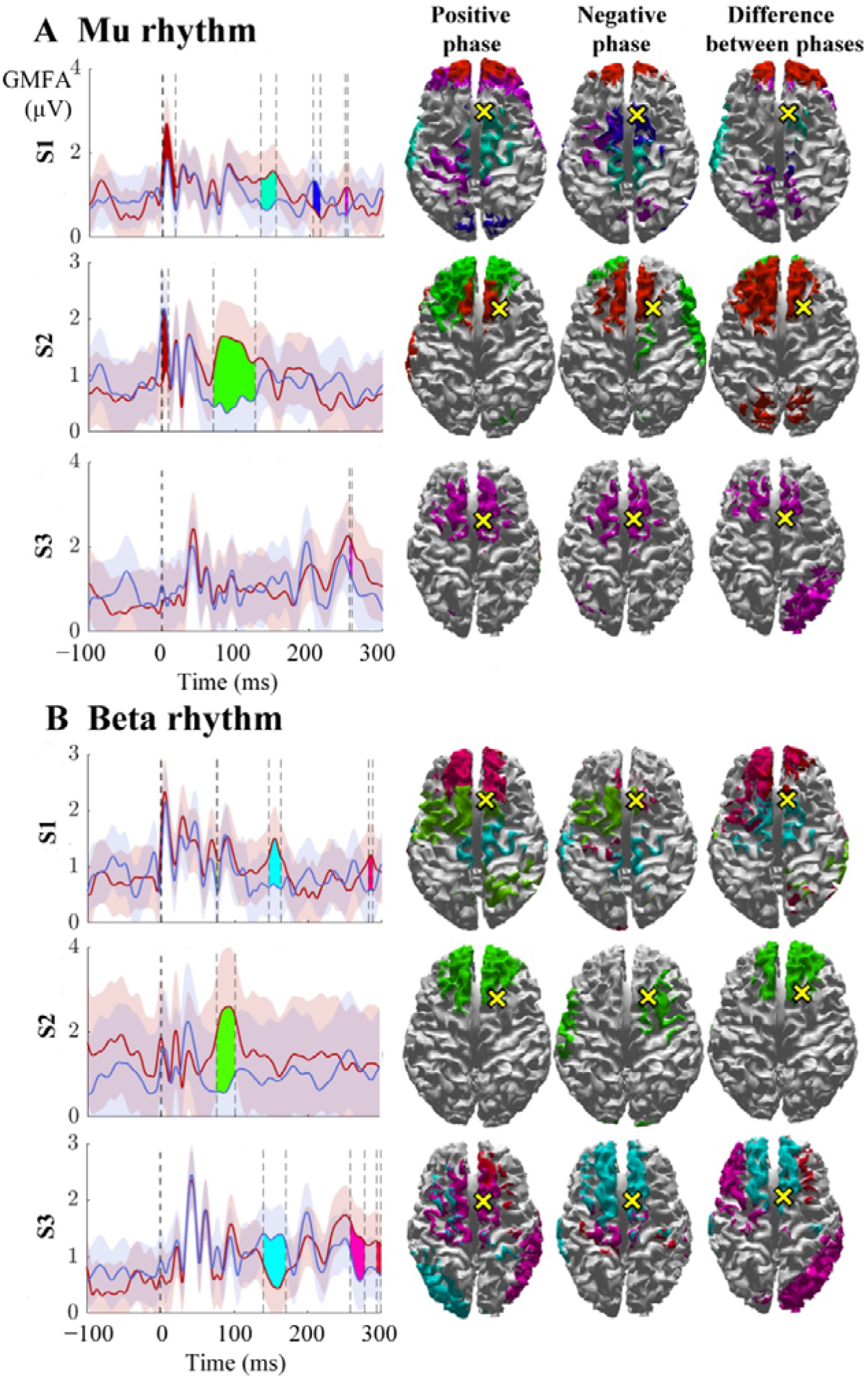
Global mean-field amplitudes (GMFA) of the positive-phase (red) and negative-phase (blue) conditions of mu (A) and beta (B) rhythms and their source estimates. The shaded areas indicate the GMFA ± the threshold for significance. Significant time intervals are marked with different colours. For each time interval, the corresponding time-averaged source estimates are shown on the right in the same colour. For each time interval, only sources stronger than 60% of the maximum amplitude are shown. The dark dashed vertical line indicates the time of the TMS pulse. The cross marks the stimulation location on pre-SMA.

### Signal propagation after M1 stimulation

In all subjects, source estimates revealed cortical activity at the stimulated site (M1) immediately after the TMS delivery, which spread to nearby frontal areas within 20 ms. By 30 ms, the activation spread more posteriorly, returning to medial frontal regions by 50 ms. After 80 ms, inter-subject variability increased, but the highest amplitudes in the source estimates remained in frontal and central areas.

The activation patterns and differences between the negative- and positive-phase classes are illustrated in Fig. 5. The mu rhythm modulated responses mainly in the frontal lobe at both early and late time points. Some differences between the classes were also found in parietal and occipital areas later than 100 ms post-stimulus. In S1, the positive-phase condition elicited stronger responses than the negative-phase condition. In S2, GMFA showed no significant TEP differences. This was likely due to larger baseline noise than in the other datasets, increasing the threshold for significance. In S3, both conditions produced larger GMFAs at different time intervals.

The differences between classes for the beta rhythm were more widespread than for the mu rhythm, although differences were most prominent in frontal and central regions. For S1, significant differences in GMFA were found only at 270–280 ms post-stimulus, when the negative-phase condition elicited stronger responses than the positive-phase condition in frontal as well as in left occipital and posterior temporal areas. In the two remaining subjects, significant differences were found prior to 50 ms as well as at 100 ms post-stimulus; S3 displayed differences also at 140 and 180 ms. For S2 and S3, differences were widespread, and spanned all four lobes.

### Signal propagation after pre-SMA stimulation

In all subjects, source estimates revealed activity at the stimulated site in the right medial frontal areas immediately after TMS. By 20–60 ms, this activation spread to anterior and to posterior frontal areas of both hemispheres. At 80 ms, source estimates also showed activation in the parietal cortex. After 100 ms, variability increased between subjects, but the strongest activity remained in frontal and parietal cortices.

The activation patterns and differences between the classes are illustrated in Fig. 6. For both rhythms, the TEP differences between the two classes were most prominent in frontal and prefrontal regions, as shown by the source estimates. For the mu rhythm, S1 displayed differences in GMFA at 0–20, 130–155, 210 and 250 ms, S2 at 0–10 and 70–130 ms, and S3 at 255 ms post-stimulus. The positive-phase condition resulted in larger absolute GMFA in five out of seven significant time intervals across subjects.

The beta rhythm modulated the responses relatively late; the earliest significant difference between the classes was found at 75 ms in S1. S1 also displayed differences centred at 155 and 280 ms post-stimulus. In S2, differences were found at 75–100 ms, with the biggest amplitude differences between the classes in the medial frontal cortex. In S3, significant differences were found at 140–170 ms and centred at 270 and 300 ms post-stimulus. The positive-phase condition resulted in larger GMFA in six out of seven significant time intervals across individuals.

## Discussion

We found that the phase of spontaneous cortical oscillations at the time of TMS appears to affect the post-stimulus effective connectivity pattern. We had proposed two hypothetical mechanisms on how the pre-stimulus phase might affect the TEPs. Based on our results, we consider the first hypothesis—that the local excitability affects only the amplitude of the cortical responses—to be less plausible than the second hypothesis. This is because the two stimulation times, at positive and negative phases of the ongoing oscillation, result in different spatial response patterns, rather than just changes in the overall amplitude.

The second hypothesis states that differences in cortical effective connectivity patterns could be explained by the local excitability state determining the preferentially activated pathways by TMS. It has been proposed that the state of the post-synaptic neural population modulates the efficacy of the synaptic transmission [46]. Such mechanisms can play a role in multiple places in the signalling cascade, determining where and when the responses differ from each other. For instance, the measured mu and beta oscillations could define the excitability of the output neurons, which are transsynaptically activated, rather than the neurons being directly excited by TMS. In that case, we would observe differences in spatial spreading with little effect on the initial responses at the stimulation site. However, in some of the cases, we observed modulation of GMFA already before the latency of 20 ms. This suggests that the phase at the stimulus onset affects neuronal activation already at the stimulated site, as it takes time for the evoked activity to spread to connected areas [70]. Nonetheless, it is possible that there were residual artefacts in the early responses, when the source estimates indicate main differences far from the stimulation site (Fig. 6A, S1). It is unlikely that artifacts depend on the oscillatory phase. However, with the limited number of trials, in some cases sporadic artefacts might distort the results. It is also possible that limitations in source estimation lead to incorrect localization of brain activity. Additionally, the poor spatial resolution of EEG makes it impossible to rule out differences in neuronal activation at the stimulated site, even when such differences are not discernible in the signal.

The correction of the phase effect (see Methods) is a crucial step in separating the evoked responses from the phase artefacts. Several studies have successfully applied this approach in eliminating spurious class differences when averaging trials with consistent phase [26, 61, 62]. As long as TMS does not significantly and consistently alter the behaviour of these background oscillations (e.g., phase or amplitude modulation), this method is robust for suppressing the background activity. However, there is a risk of overcorrecting the data, although such an error is smaller than the error introduced by the phase effect [26].

In a strict sense, our statistical analysis of GMFA did not include a multiple comparisons correction, which increases the risk of reporting false positives. To address this, we tested the sensitivity of the method by randomly assigning trials to each class, which produced none or markedly less significant differences than the real classes. However, future studies should be conducted to confirm our observations with more confidence.

Patterns of effective connectivity are known to be subject-specific [71]. Therefore, it is not surprising that the spreading of TMS-evoked activity as a function of the oscillation phase at pulse time differed across the subjects. This variability might be due to differences in the connections between the stimulation site and the connected ones or in the exact stimulus location and the intracortical fiber pathways that are activated. In addition to the possible inter-subject variability in the probed brain connectivity, the phase estimation is a plausible source of further variability. The C3 and F2 channels do not measure exactly the same cortical sources in different subjects. For instance, details in the cortical folding significantly influence the EEG sensitivity patterns.

Our observations that the spontaneous oscillations modulate effective connectivity mainly within 80–170 ms after the stimulus are in line with Fehér et al. [72], who reported that the phase of transcranial alternating current stimulation modulates cortical signal propagation. They observed significant phase-dependent modulation of TMS-evoked responses starting at 40–60 ms and lasting until 200 ms post-stimulus when stimulating the dorsolateral prefrontal cortex.

Finally, this study highlights the importance of brain state on signal propagation after TMS. This can be a crucial aspect for repetitive TMS (rTMS) commonly used for therapy. rTMS is known to modulate the excitability and connectivity of stimulated areas, contributing to the treatment of, e.g., major depressive disorder (MDD) and movement disorders [73]. The pre-stimulus mu rhythm at high and low excitability states affects the rTMS effect in M1 on MEPs [10]. Similarly, alpha-synchronized rTMS has shown potential in MDD therapy [30]. The efficacy of rTMS strategies for treatment and rehabilitation of various neurological and psychiatric disorders may be connected to the preferred signal pathways based on the excitability states right before the delivery of rTMS. Further studies are needed to investigate the effect at different targets and different intensities, as well as the role of the propagation pattern in the treatment outcome. This can be crucial in deciding which rTMS paradigm is most beneficial in different disorders at the single patient level [74].

## Conclusions

Delivering TMS during distinct cortical excitability states resulted in different signal propagation patterns, suggesting that effective connectivity is phase-dependent. The phase of local cortical oscillations modulates not only the local excitability, but also the overall activation of different overlapping neuronal populations. Our findings open new avenues for further research and may be crucial for the success of future brain-state-dependent and closed-loop stimulation paradigms in patient care.

## Acknowledgments

This project has received funding from the European Research Council (ERC Synergy) under the European Union’s Horizon 2020 research and innovation programme (ConnectToBrain; grant agreement No. 810377), the Academy of Finland (Decisions No. 294625, 321631 and 327326), Jane and Aatos Erkko Foundation, the Finnish Cultural Foundation, and the Instrumentarium Science Foundation. This research has also received funding from the European Union’s Horizon 2020 Framework Programme for Research and Innovation under the Specific Grant Agreement No. 945539 (Human Brain Project SGA3 to M.R.), by Fondazione Regionale per la Ricerca Biomedica (Regione Lombardia), Project ERAPERMED2019-101, GA 779282 (to M.R.), and from the Tiny Blue Dot Foundation (to M.R. and M.F.).

